## Supplemental Information and Figures for "Molecular Consequences of Peripheral Influenza A Infection on Cell Populations in the Murine Hypothalamus"

**Supplementary Information**

**Supplementary Tables**

- **Supplementary Table 1 (Supp.table01_CellCounts)**
  Number of cells in different cell populations across different cluster levels
  - **Sheet QC counts**: The dataset underwent different quality control steps. This tables contains the amount of cells per animal at the different control and filter steps.
  - **Sheet Cell.counts_celltype.lev1**: Table contains the amount of cells for the different sequenced animals in neuronal or non-neuronal cells.
  - **Sheet Cell.counts_celltype.lev2**:This table contains the amount of cells per anima across all GABAergic, glutamatergic or non-neuronal cells.
  - **Sheet Cell.counts_cluster.id**: This table shows the number of cells per animal for the different identified cell clusters.

- **Supplementary Table 2 (Supp.table02_PredictionsPercentPerCellType_forPredScoreGreate50Perc)**
  Results from cell label transfer from different datasets (see Methods) on cell cluster level. Containing information about the dataset, reference data ID, number of total cells, number of cells that fitted the prediction score criteria (score > 0.5) and percent of cells in cell type tagged with selected prediction score cut-off.
  - **Sheet Glut**: Results from the cell label transfer for glutamatergic clusters.
  - **Sheet Gaba**: Results from the cell label transfer for GABAergic clusters.
  - **Sheet NonN**: Results from the cell label transfer for non-neuronal clusters.

- **Supplementary Table 3 (Supp.table03_Cellcluster_annotations)**
  Predicted cell cluster annotations based on cell label transfer.
  - Sheet Selection criteria: Describes the criteria for transferring cell labels to the identified clusters.
  - **Sheet NonN**: Cell labels for non-neuronal clusters.
  - **Sheet Glut**: Cell labels for glutamatergic clusters.
  - **Sheet GABA**: Cell labels for GABAergic clusters.

- **Supplementary Table 4 (Supp.table04_MarkerGenes)**
  Marker genes identified using a receiver operating characteristic (ROC) method.
  - **Sheet GABA**: List of marker genes for GABAergic clusters.
  - **Sheet Glut**: List of marker genes for glutamatergic clusters.
  - **Sheet NonN:** List of marker genes for non-neuronal clusters.

- **Supplementary Table 5 (Supp.table05_DEGs)**
  Results from pseudobulb differential gene expression analysis. The analysis was performed 3 times on different cell type levels.
  - **Sheet 3dpi - DEG - across all cells**: Differential expressed genes comparing all control cells vs all cells at 3dpi.
  - **Sheet 7dpi - DEG - across all cells**: Differential expressed genes comparing all control cells vs all cells at 7dpi.
  - **Sheet 23dpi - DEG - across all cells**: Differential expressed genes comparing all control cells vs all cells at 23dpi.

- **Sheet GABA - DEG - across celltype**: Differential expressed genes from a pseudo-bulk analysis of GABAergic cells.
- **Sheet Glut - DEG - across celltype**: Differential expressed genes from a pseudo-bulk analysis of glutamatergic cells.
- **Sheet NonN - DEG - across celltype**: Differential expressed genes from a pseudo-bulk analysis of non-neuronal cells.
- **Sheet GABA - DEG - across cellcluster**: Differential expressed genes from a pseudo-bulk analysis on GABAergic cell cluster levels.
- **Sheet Glut - DEG - across cellcluster**: Differential expressed genes from a pseudo-bulk analysis on glutamatergic cell cluster levels.
- **Sheet NonN - DEG - across cellcluster**: Differential expressed genes from a pseudo-bulk analysis on non-neuronal cell cluster levels.

- **Supplementary Table 6 (Supp.table06_DEGs_SubAnalysis_Neurons)**
  Results from pseudo-bulk differential gene expression analysis with less restrictive filter criteria on selected neuronal cell clusters (see Methods).
  - **Sheet 3dpi**: Analysis results for comparison Control vs 3dpi.
  - **Sheet 7dpi**: Analysis results for comparison Control vs 7dpi.
  - **Sheet 23dpi**: Analysis results for comparison Control vs 23dpi.

- **Supplementary Table 7 (Supp.table07_ReactomeEnrichment)**
  Results from Reactome pathway enrichment analysis.

- **Supplementary Table 8 (Supp.table08_GOenrichment)**
  Results from Gene Ontology (GO) enrichment analysis.

- **Supplementary Table 9 (Supp.table09_Cacoa_geneprograms)**
  Gene programs from cluster free expression shift analysis from Cacoa package. Analysis was performed for Control against the 3 different time points separately.
  - **Sheet Program scores UMAP embeddings**: Contains the Adjusted z-scores from the identified genes programs at the different time points in all cells.
  - **Sheet 3 dpi - gene programs**: Contains gene names, sim scores and loadings (for more information see (Petukhov et al., 2022)) for 9 different gene programs identified comparing cells from control samples against cells at 3 dpi.
  - **Sheet 7 dpi - gene programs**: Contains gene names, sim scores and loadings (for more information see (Petukhov et al., 2022)) for 9 different gene programs identified comparing cells from control samples against cells at 7 dpi.
  - **Sheet 23 dpi - gene programs**: Contains gene names, sim scores and loadings (for more information see (Petukhov et al., 2022)) for 8 different gene programs identified comparing cells from control samples against cells at 23 dpi.

### Supplementary Figures

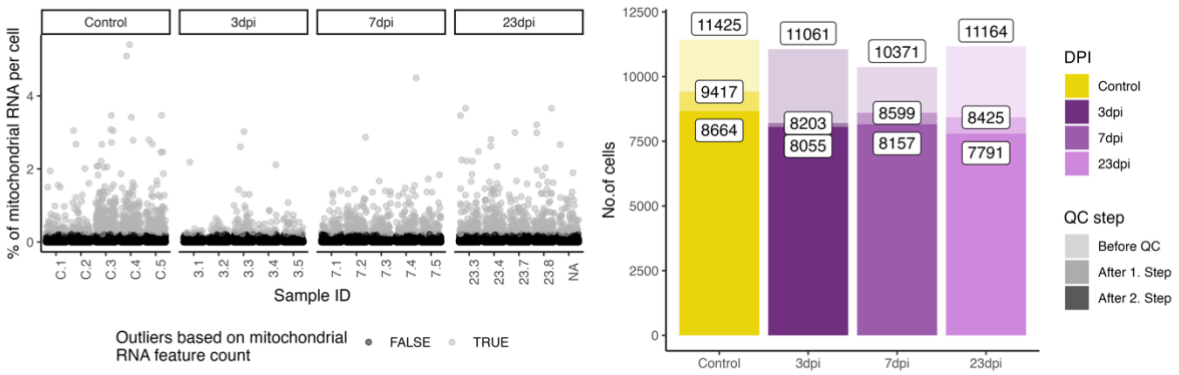


Supplementary Figure 1 **Quality Control plots for the generated snRNA-seq dataset. A.** Percentages of mitochondrial RNA in the different samples. Light gray dots were categorised as outliers and removed form the downstream analysis. **B.** Cell counts per time point after different filter steps.

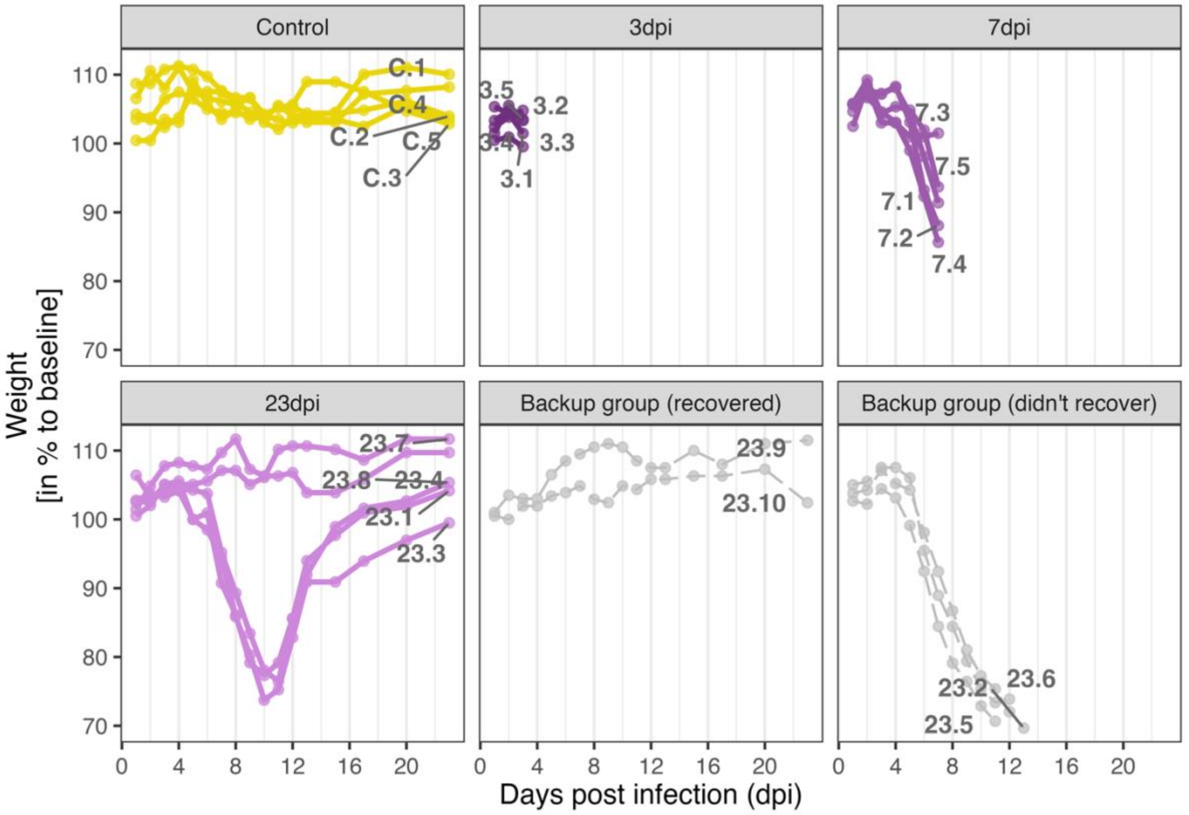


Supplementary Figure 2 **Weight curves depicting the weight-loss due to H1N1 pdm09 infection in the different individuals.** Weight curves are group by time point samples. Grey coloured animals were not used for snRNA-sequencing.

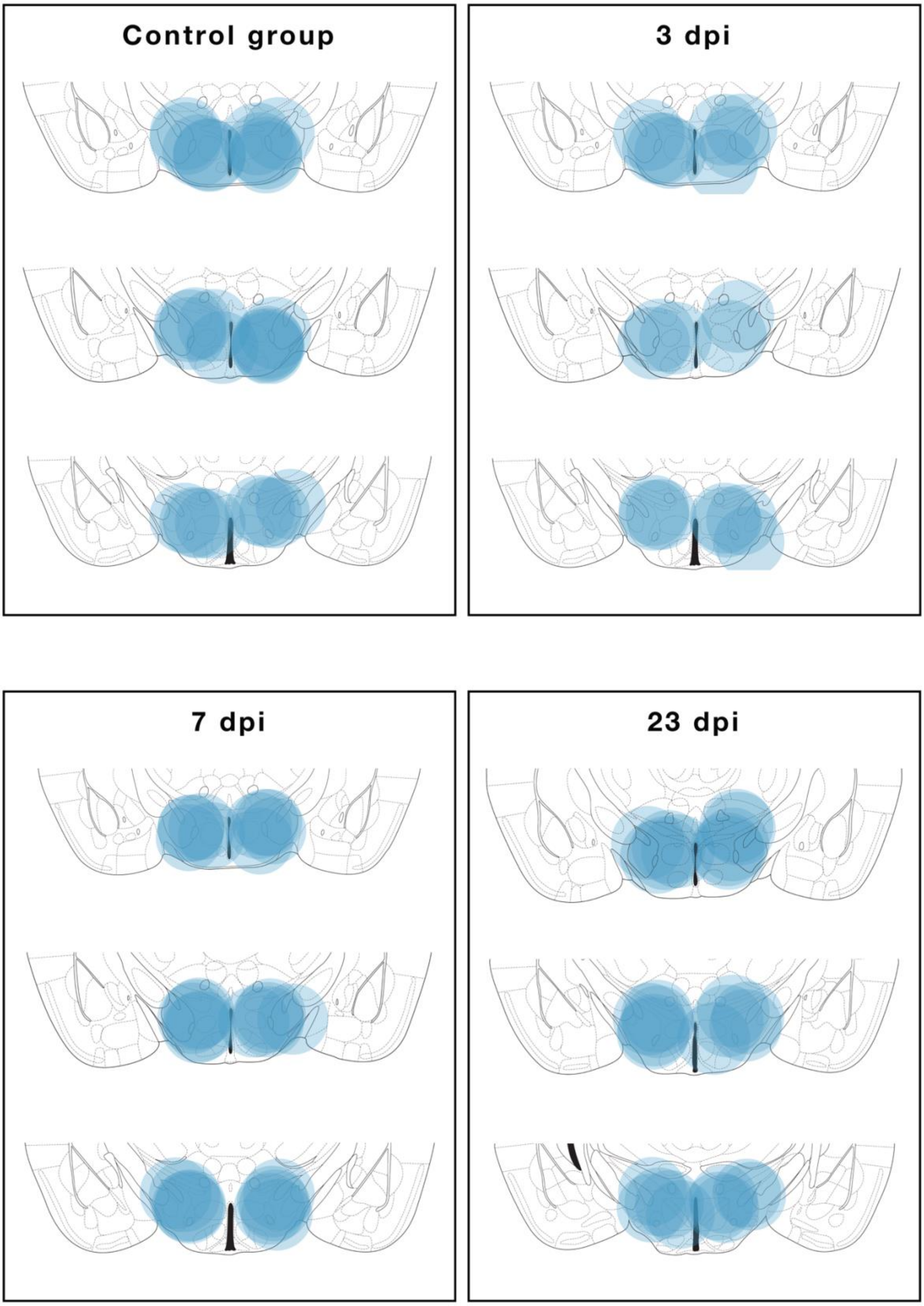


Supplementary Figure 3 **Punching location.** Location of 4 mm punches for RNA extraction and snRNA-seq in the hypothalamus in the different sampling groups.

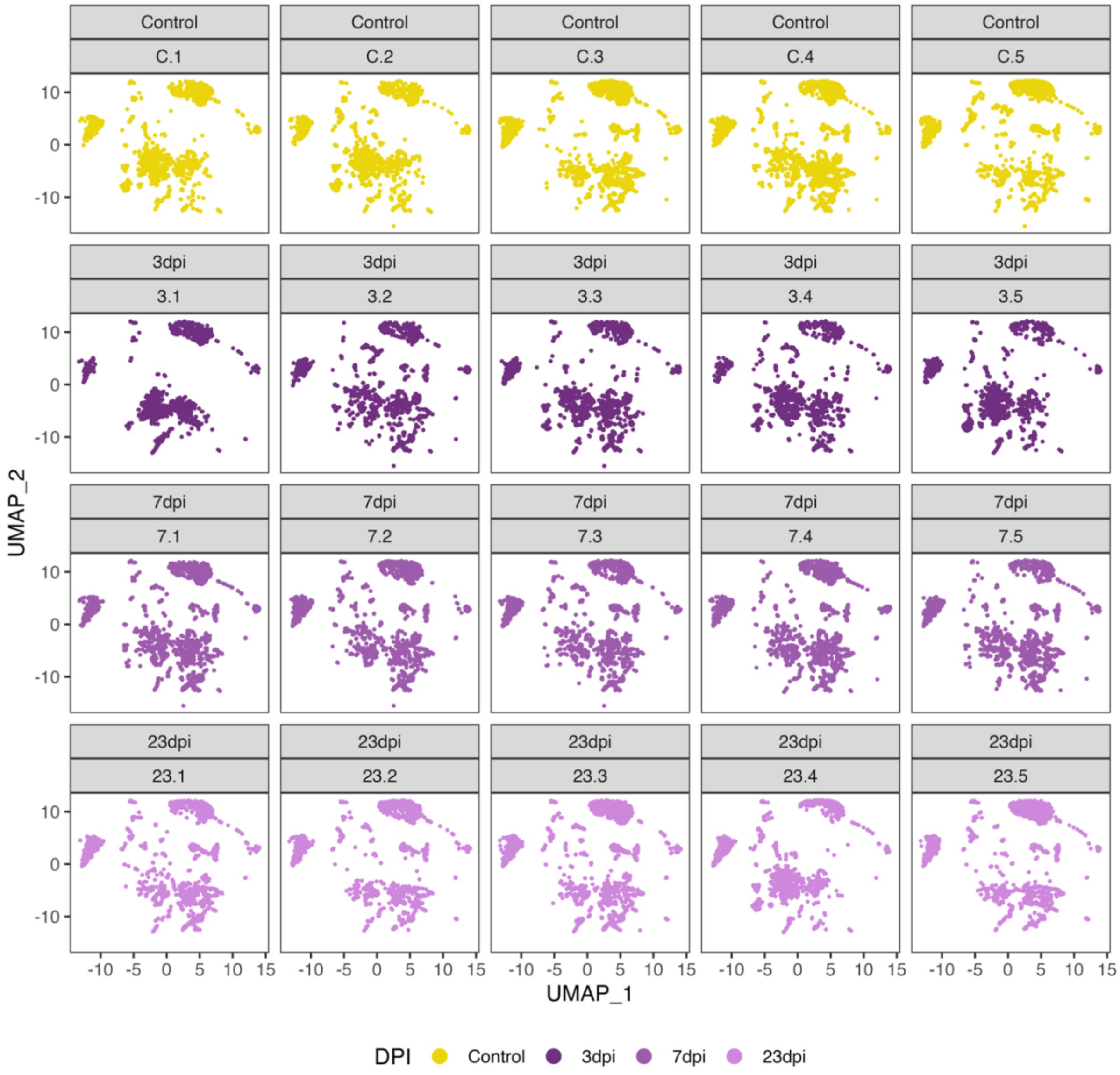


Supplementary Figure 4 **UMAP embeddings per hash-tagged sample.** UMAP embeddings show the distribution of cells across the different cell types of the individual hash-tagged samples. Colours depict and shadings depict the time points.

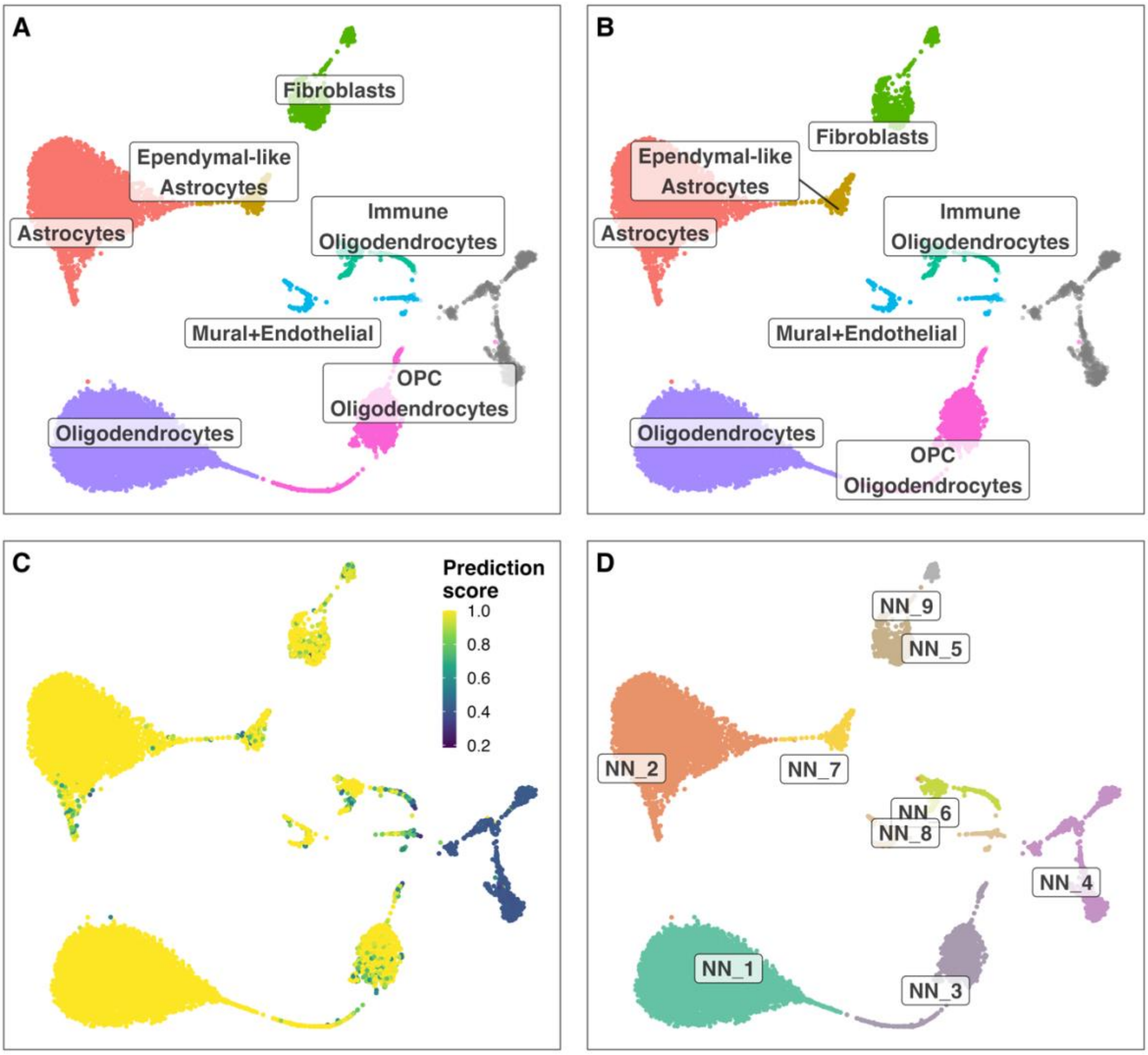


F

| Cluster ID | Predicted ID | Predicted name | # Total cells | # Predic. cells | Mean predic. score | % Predic. cells |
|---|---|---|---|---|---|---|
| NN_1 | C25-19 | Oligodendrocytes | 5023 | 5022 | 1.00 | 99.98 |
| NN_2 | C25-18 | Astrocytes | 4542 | 4510 | 1.00 | 99.30 |
| NN_3 | C25-20 | OPC | 1106 | 934 | 0.97 | 84.45 |
| NN_3 | C25-19 | Oligodendrocytes | 1106 | 149 | 0.93 | 13.47 |
| NN_5 | C25-25 | Fibroblasts | 492 | 457 | 0.97 | 92.89 |
| NN_6 | C25-21 | Immune | 339 | 236 | 0.92 | 69.62 |
| NN_6 | C25-19 | Oligodendrocytes | 339 | 48 | 0.84 | 14.16 |
| NN_7 | C25-17 | Ependymal-like | 321 | 268 | 0.94 | 83.49 |
| NN_7 | C25-18 | Astrocytes | 321 | 33 | 0.88 | 10.28 |
| NN_8 | C25-24 | Mural+Endothelial | 219 | 166 | 0.95 | 75.80 |
| NN_9 | C25-25 | Fibroblasts | 134 | 117 | 0.92 | 87.31 |

Supplementary Figure 5 **Label transfer of cell-type labels from the HypoMap (Steuernagel et al., 2022) annotations (C7) to the Non-neuronal cell cluster here identified. A/B.** Shows the cell label transfer for HypoMap cell-type level C7. **C.** Depicts prediction scores. **D.** Original cell-clusters and labels identified in the here presented study. F. Detailed overview of predicted names for the different identified cell clusters.

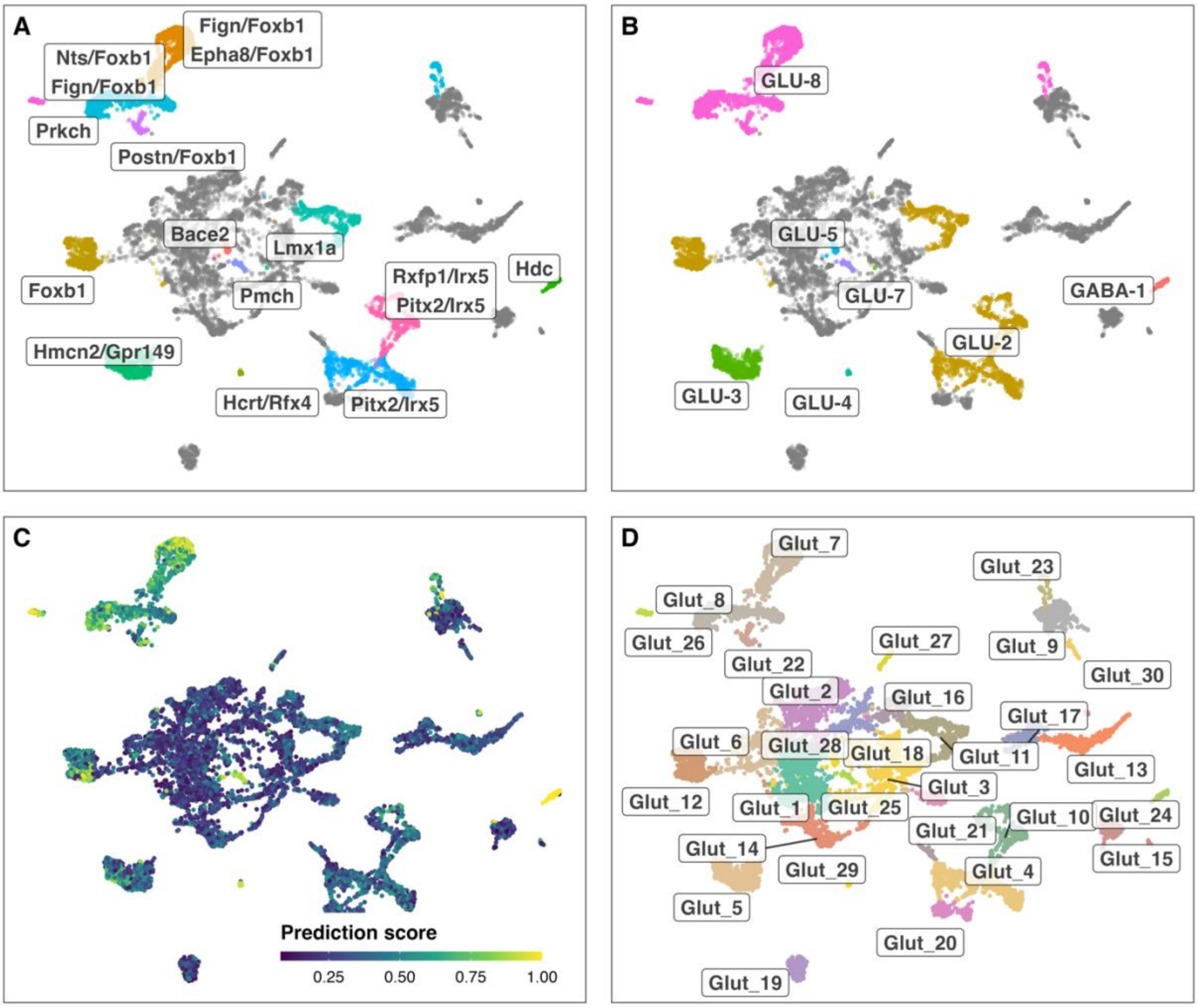


F

| Cluster ID | Predicted ID | Predicted name | Predic. Cell ID | Markers | # Total cells | # Predic. cells | Mean predic. score | % Predic. cells |
|---|---|---|---|---|---|---|---|---|
| Glut_10 | C185-6 | Rxfp1.Irx5.GLU-2 | GLU-2 | Rxfp1/Irx5 | 520 | 73 | 0.62 | 14.04 |
| Glut_10 | C185-4 | Pitx2.Irx5.GLU-2 | GLU-2 | Pitx2/Irx5 | 520 | 62 | 0.58 | 11.92 |
| Glut_11 | C185-20 | Lmx1a.GLU-2 | GLU-2 | Lmx1a | 476 | 83 | 0.56 | 17.44 |
| Glut_12 | C185-19 | Foxb1.GLU-2 | GLU-2 | Foxb1 | 476 | 241 | 0.78 | 50.63 |
| Glut_22 | C185-58 | Postn.Foxb1.GLU-8 | GLU-8 | Postn/Foxb1 | 100 | 43 | 0.75 | 43.00 |
| Glut_23 | C185-57 | Nts.Foxb1.GLU-8 | GLU-8 | Nts/Foxb1 | 98 | 30 | 0.68 | 30.61 |
| Glut_23 | C185-59 | Fign.Foxb1.GLU-8 | GLU-8 | Fign/Foxb1 | 98 | 27 | 0.71 | 27.55 |
| Glut_24 | C185-102 | Hdc.GABA-1 | GABA-1 | Hdc | 94 | 91 | 0.99 | 96.81 |
| Glut_25 | C185-56 | Pmch.GLU-7 | GLU-7 | Pmch | 85 | 80 | 0.86 | 94.12 |
| Glut_26 | C185-61 | Prkch.GLU-8 | GLU-8 | Prkch | 80 | 76 | 0.93 | 95.00 |
| Glut_28 | C185-52 | Bace2.GLU-5 | GLU-5 | Bace2 | 64 | 26 | 0.64 | 40.62 |
| Glut_29 | C185-47 | Hcrt.Rfx4.GLU-4 | GLU-4 | Hcrt/Rfx4 | 38 | 35 | 0.92 | 92.11 |
| Glut_4 | C185-4 | Pitx2.Irx5.GLU-2 | GLU-2 | Pitx2/Irx5 | 744 | 84 | 0.57 | 11.29 |
| Glut_5 | C185-30 | Hmcn2.Gpr149.GLU-3 | GLU-3 | Hmcn2/Gpr149 | 694 | 148 | 0.67 | 21.33 |
| Glut_7 | C185-59 | Fign.Foxb1.GLU-8 | GLU-8 | Fign/Foxb1 | 625 | 386 | 0.81 | 61.76 |
| Glut_7 | C185-60 | Epha8.Foxb1.GLU-8 | GLU-8 | Epha8/Foxb1 | 625 | 164 | 0.65 | 26.24 |
| Glut_8 | C185-57 | Nts.Foxb1.GLU-8 | GLU-8 | Nts/Foxb1 | 604 | 305 | 0.71 | 50.50 |
| Glut_8 | C185-59 | Fign.Foxb1.GLU-8 | GLU-8 | Fign/Foxb1 | 604 | 75 | 0.63 | 12.42 |

Supplementary Figure 6 **Label transfer of cell-type labels from the HypoMap (Steuernagel et al., 2022) annotations (C285) to glutamatergic cell cluster here identified. A/B.** Shows the cell label transfer for HypoMap cell-type level C285 (_named). **C.** Depicts prediction scores. **D.** Original cell-clusters and labels identified in the here presented study. **F.** Detailed overview of predicted names for the different identified cell clusters.

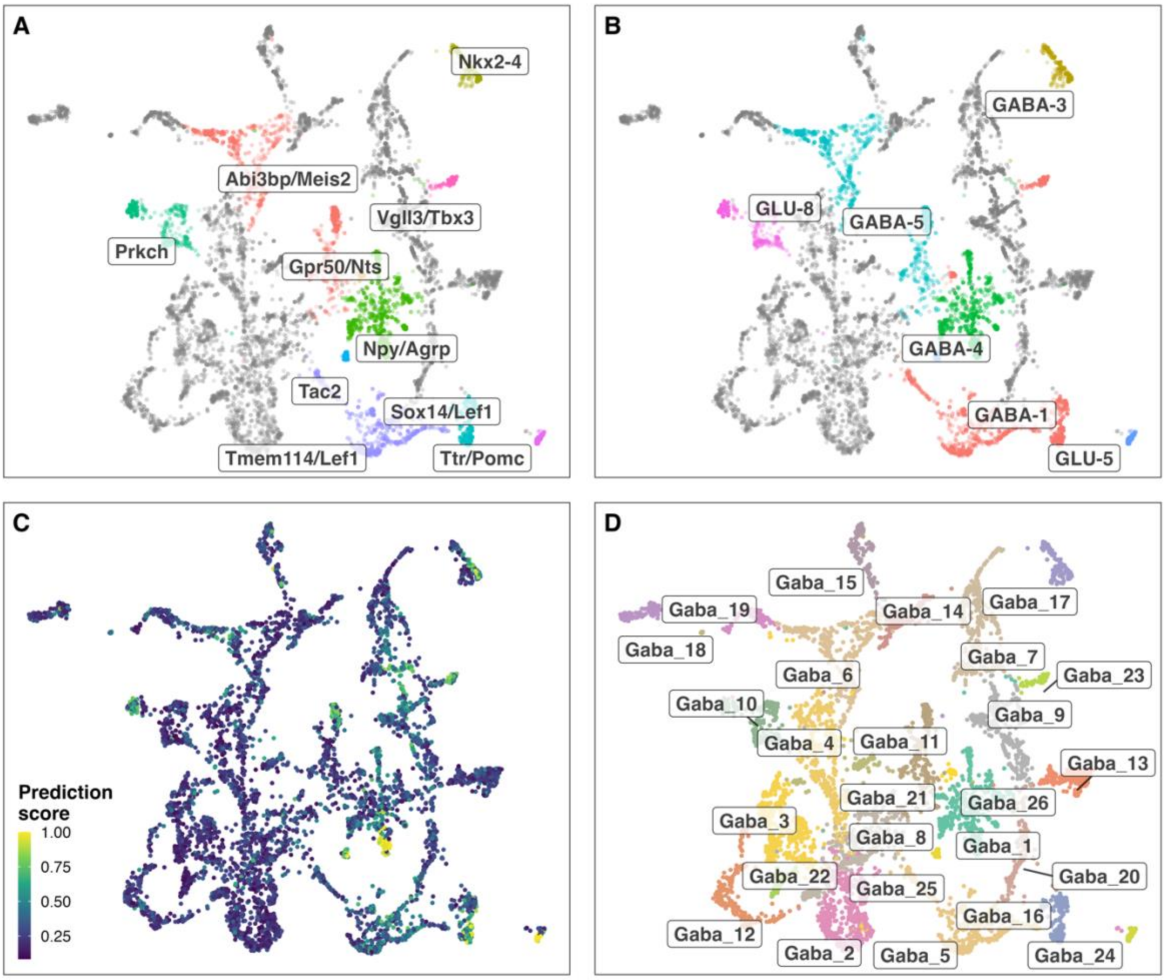


F

| Cluster ID | Predicted ID | Predicted name | Predic. Cell ID | Markers | # Total cells | # Predic. cells | Mean predic. score | % Predic. cells |
|---|---|---|---|---|---|---|---|---|
| Gaba_1 | C185-115 | Npy.Agrp.GABA-4 | GABA-4 | Npy/Agrp | 486 | 64 | 0.96 | 13.17 |
| Gaba_10 | C185-61 | Prkch.GLU-8 | GLU-8 | Prkch | 265 | 38 | 0.73 | 14.34 |
| Gaba_11 | C185-121 | Abi3bp.Meis2.GABA-5 | GABA-5 | Abi3bp/Meis2 | 250 | 60 | 0.71 | 24.00 |
| Gaba_16 | C185-88 | Sox14.Lef1.GABA-1 | GABA-1 | Sox14/Lef1 | 161 | 76 | 0.77 | 47.20 |
| Gaba_17 | C185-113 | Nkx2-4.GABA-3 | GABA-3 | Nkx2-4 | 144 | 39 | 0.70 | 27.08 |
| Gaba_23 | C185-103 | Vgll3.Tbx3.GABA-1 | GABA-1 | Vgll3/Tbx3 | 68 | 35 | 0.74 | 51.47 |
| Gaba_24 | C185-50 | Ttr.Pomc.GLU-5 | GLU-5 | Ttr/Pomc | 62 | 36 | 0.95 | 58.06 |
| Gaba_25 | C185-51 | Tac2.GLU-5 | GLU-5 | Tac2 | 41 | 37 | 0.77 | 90.24 |
| Gaba_26 | C185-110 | Gpr50.Nts.GABA-1 | GABA-1 | Gpr50/Nts | 29 | 8 | 0.62 | 27.59 |
| Gaba_5 | C185-86 | Tmem114.Lef1.GABA-1 | GABA-1 | Tmem114/Lef1 | 394 | 46 | 0.62 | 11.68 |
| Gaba_6 | C185-121 | Abi3bp.Meis2.GABA-5 | GABA-5 | Abi3bp/Meis2 | 378 | 40 | 0.66 | 10.58 |

Supplementary Figure 7 **Label transfer of cell-type labels from the HypoMap (Steuernagel et al., 2022) annotations (C285) to GABAergic cell cluster here identified. A/B.** Shows the cell label transfer for HypoMap cell-type level C285 (_named). **C.** Depicts prediction scores. **D.** Original cell-clusters and labels identified in the here presented study. **F.** Detailed overview of predicted names for the different identified cell clusters.

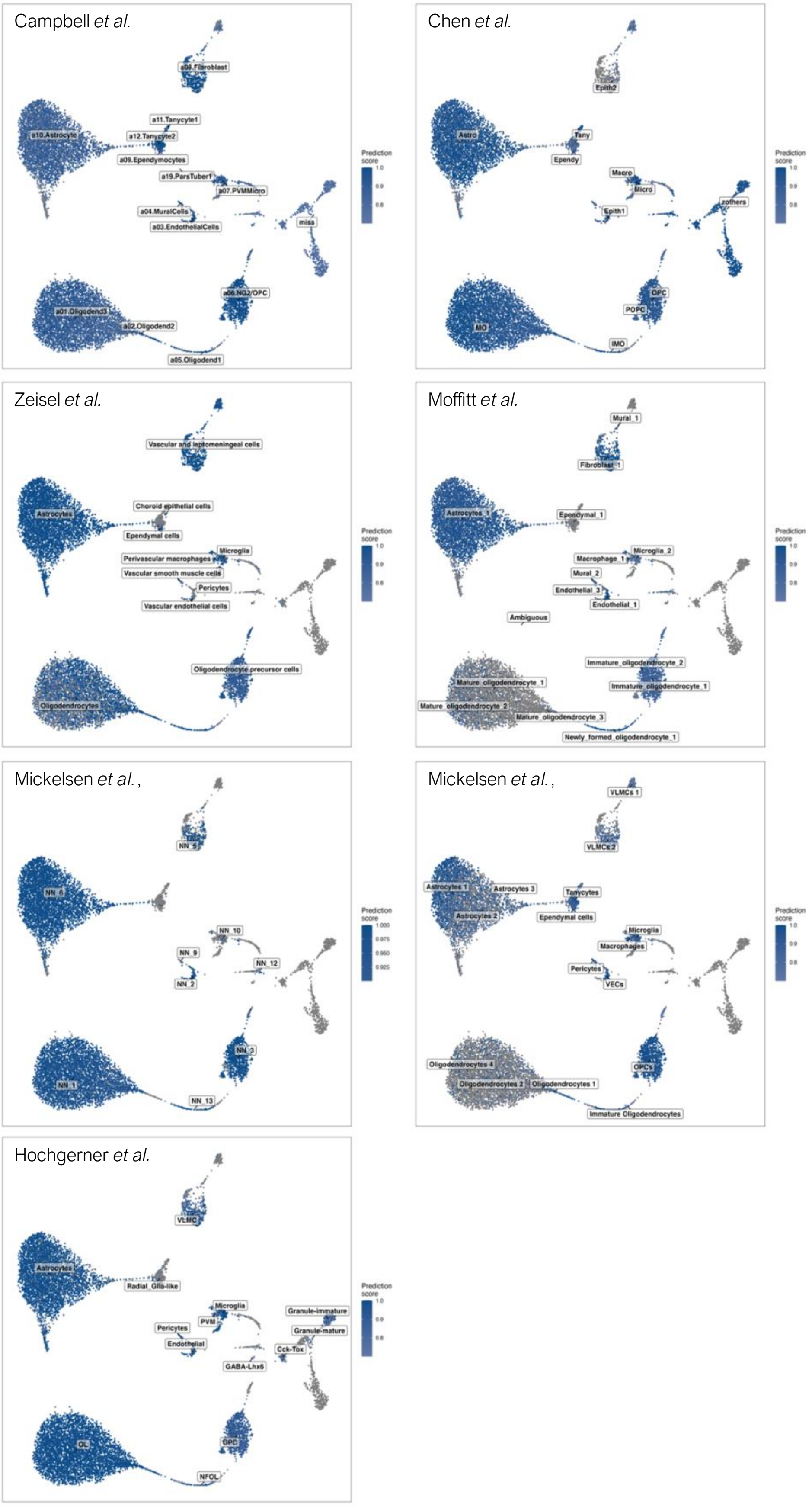


Supplementary Figure 8 **Cell-type label transfer for non-neuronal cells.** Depicting the prediction scores and potential cell-type labels in non-neuronal cells from different published datasets (Campbell et al., 2017; Chen et al., 2017; Hochgerner et al., 2018; Mickelsen et al., 2019; Mickelsen et al., 2020; Moffitt et al., 2018; Zeisel et al., 2018).

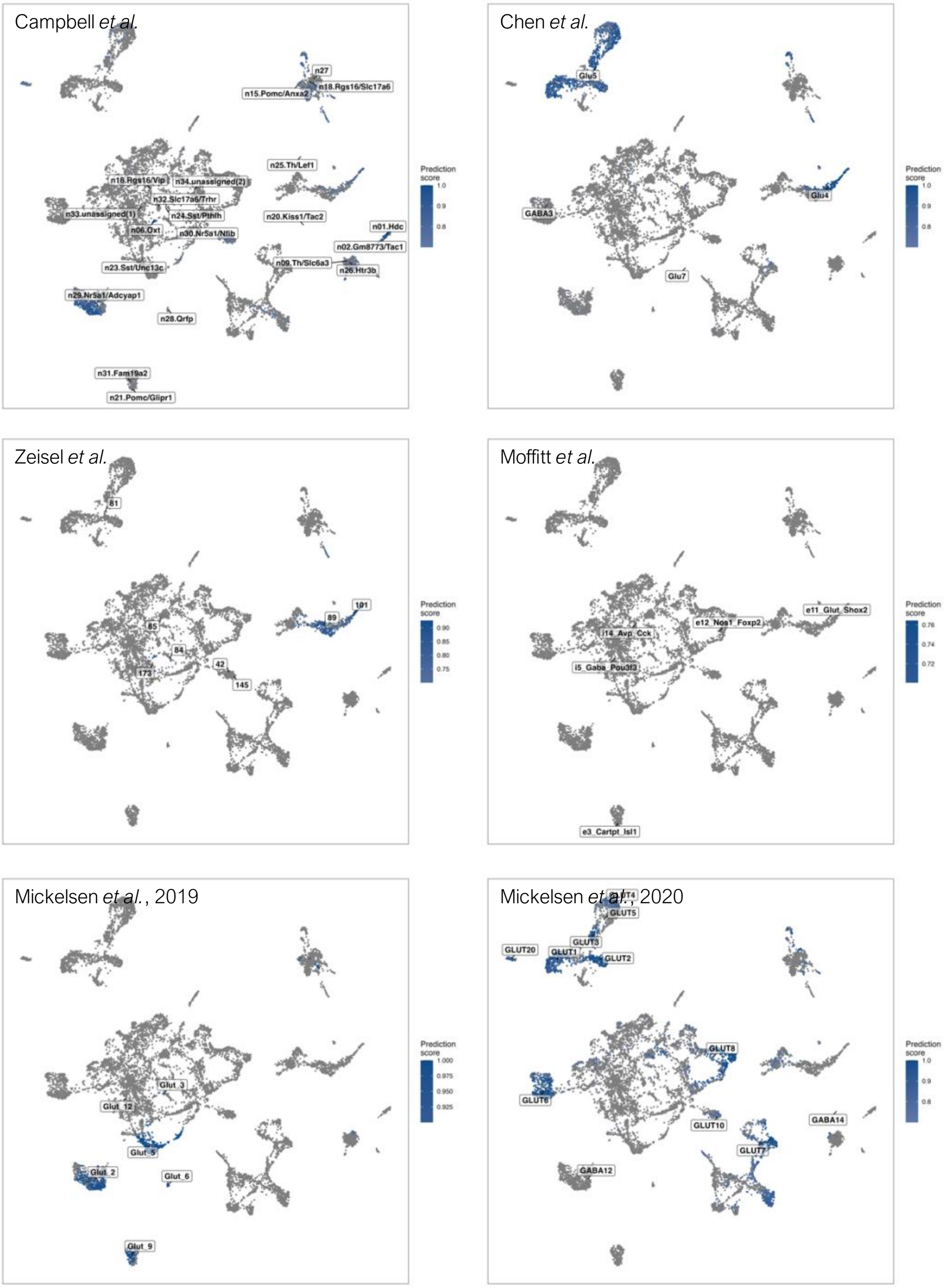


Supplementary Figure 9 **Cell-type label transfer for glutamatergic cells.** Depicting the prediction scores and potential cell-type labels in non-neuronal cells from different published datasets (Campbell et al., 2017; Chen et al., 2017; Mickelsen et al., 2019; Mickelsen et al., 2020; Moffitt et al., 2018; Zeisel et al., 2018).

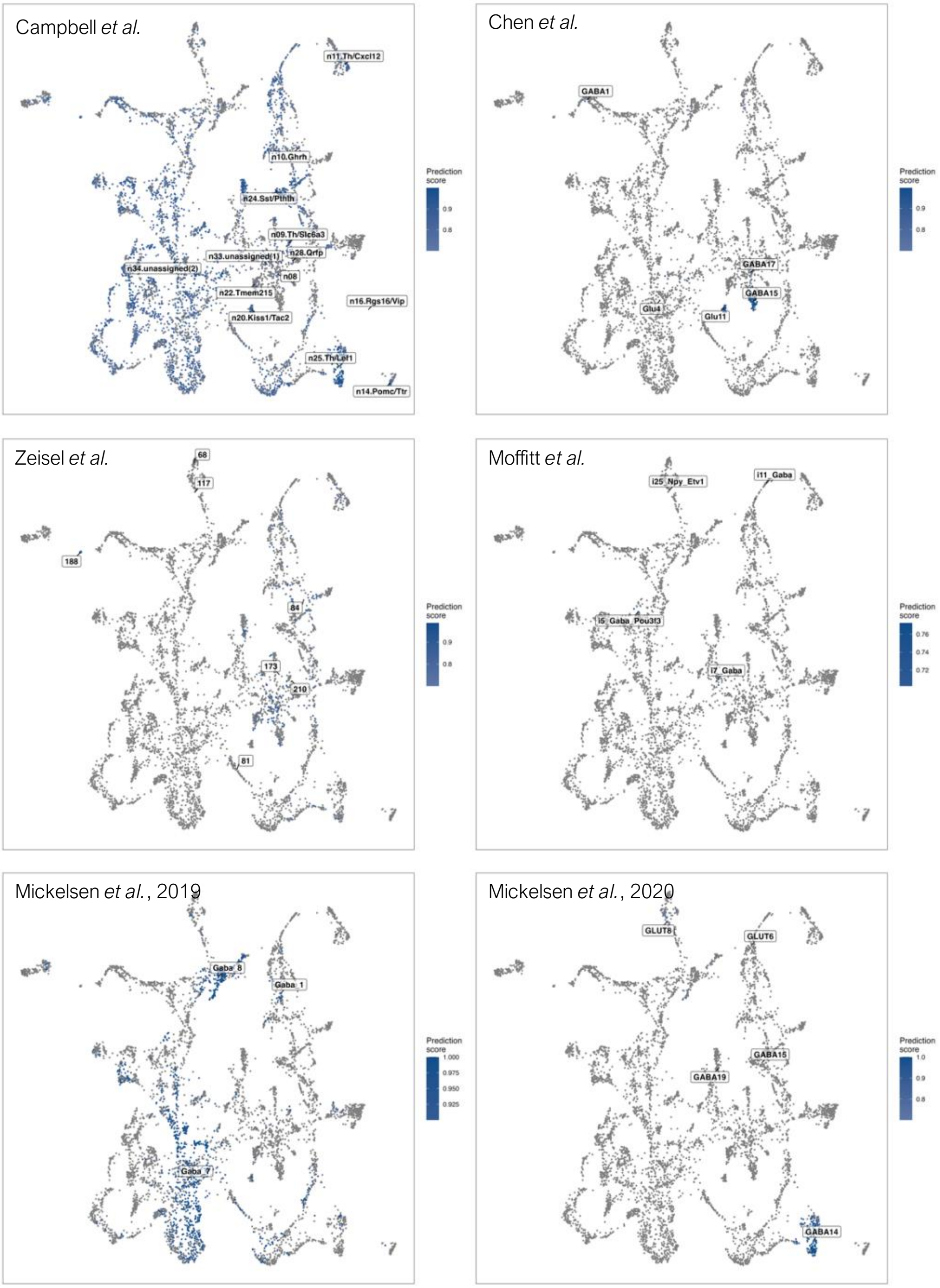


Supplementary Figure 10 **Cell-type label transfer for glutamatergic cells.** Depicting the prediction scores and potential cell-type labels in non-neuronal cells from different published datasets (Campbell et al., 2017; Chen et al., 2017; Mickelsen et al., 2019; Mickelsen et al., 2020; Moffitt et al., 2018; Zeisel et al., 2018).

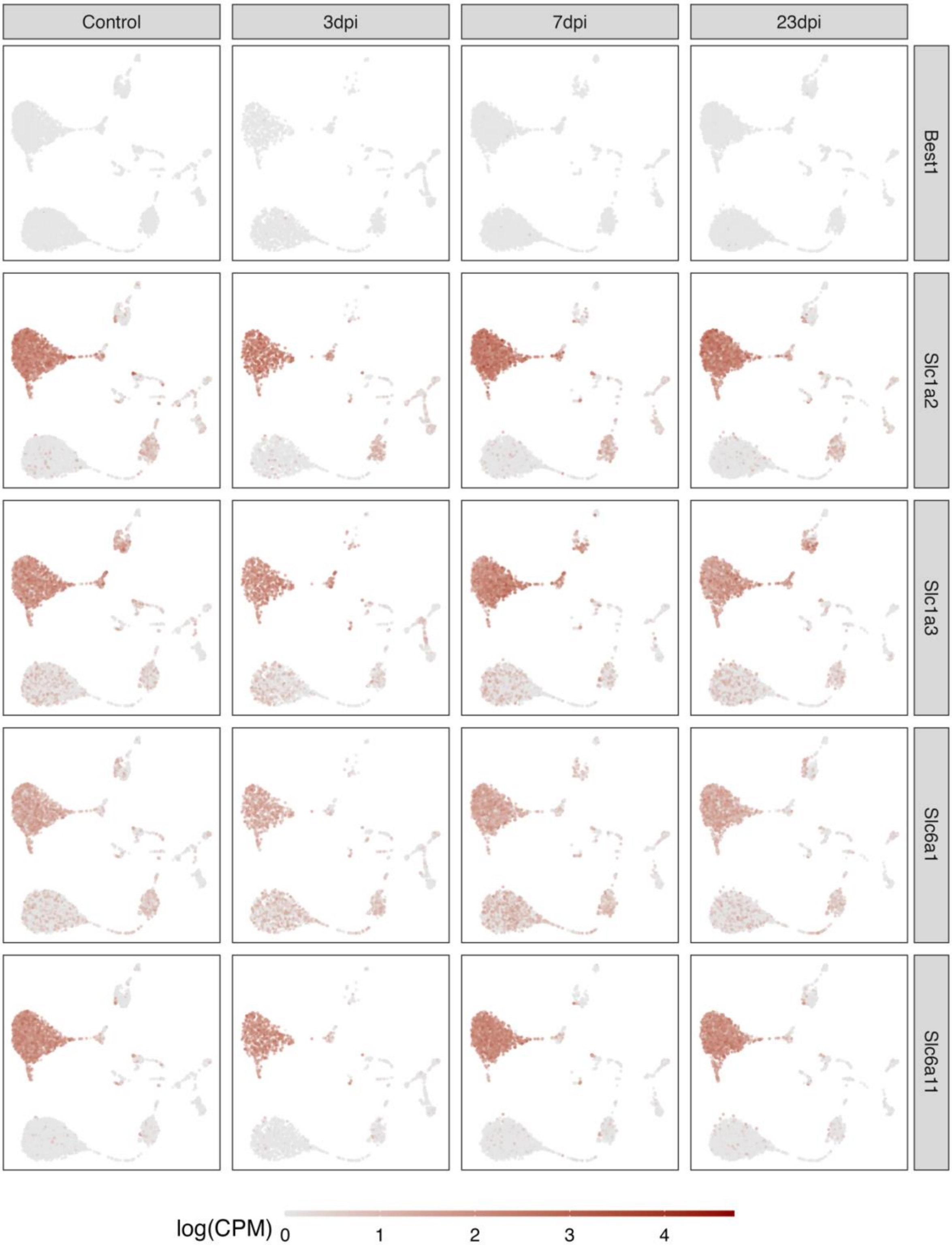


Supplementary Figure 11 **Expression of different GABAergic and glutamatergic transporters.** Depicted are normalized expression levels in non-neuronal cells of different GABAergic and glutamatergic transporters.

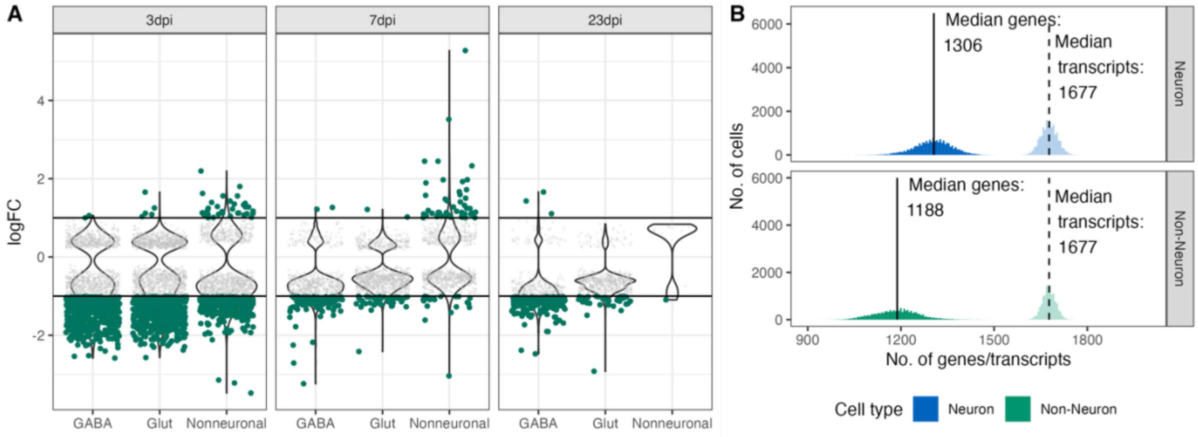


Supplementary Figure 12 **DEG analysis in down-sampled dataset. A.** Scatter- and Violin-plot showing the results of a differential gene expression analysis comparing infected groups against control groups on all Glutamatergic, GABAergic and Non-neuronal cells on a down sampled data set. **B**. Histogram showing the distribution of features (genes) and transcripts per cell in a down0sampled dataset (down-sampled to 1600 transcripts per cell).

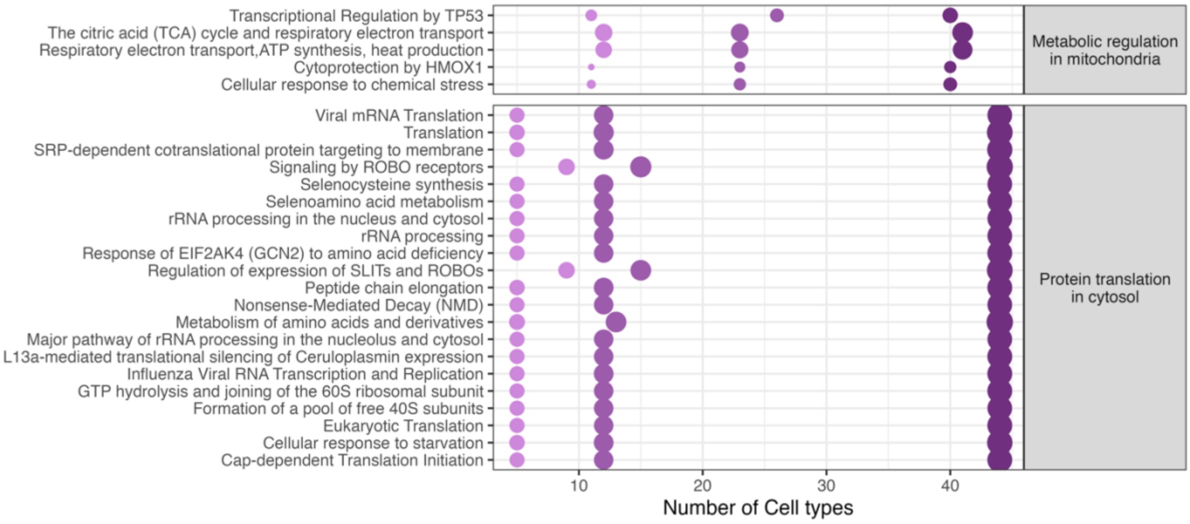


Supplementary Figure 13 **Reactome pathway enrichment of differentially expressed genes in neurons.** Dotplot shows significantly enriched Reachtome pathways in neuronal cell popluations across different timepoints. Gradients of purple depict reflect the time points of analysis. 3 dpi (dark purple) to 23dpi (light purple).

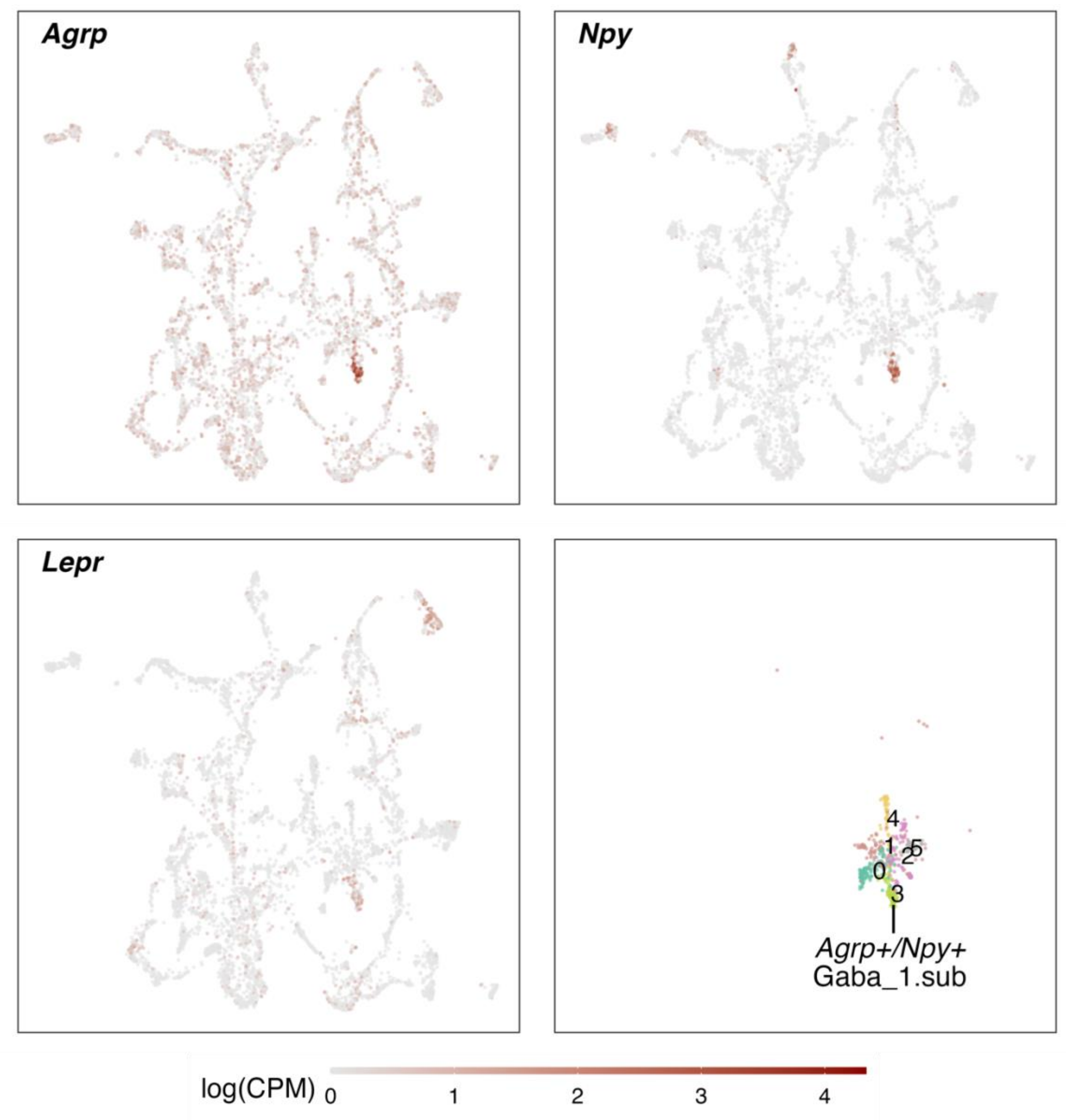


Supplementary Figure 14 **Identification of an *Agrp+/Npy+* neuron cluster.** *Agrp*+/*Npy*+ neurons were identified as a subcluster of the GABA_1 neuron population. Depicted are the normalized expression levels of marker genes *Agrp*, *Npy* and *Lepr*, showing their distinct expression in a cell cluster within the GABA_1 cluster. An additional cluster analysis of the GABA_1 cluster, identified them in the subcluster 3 (lower left).

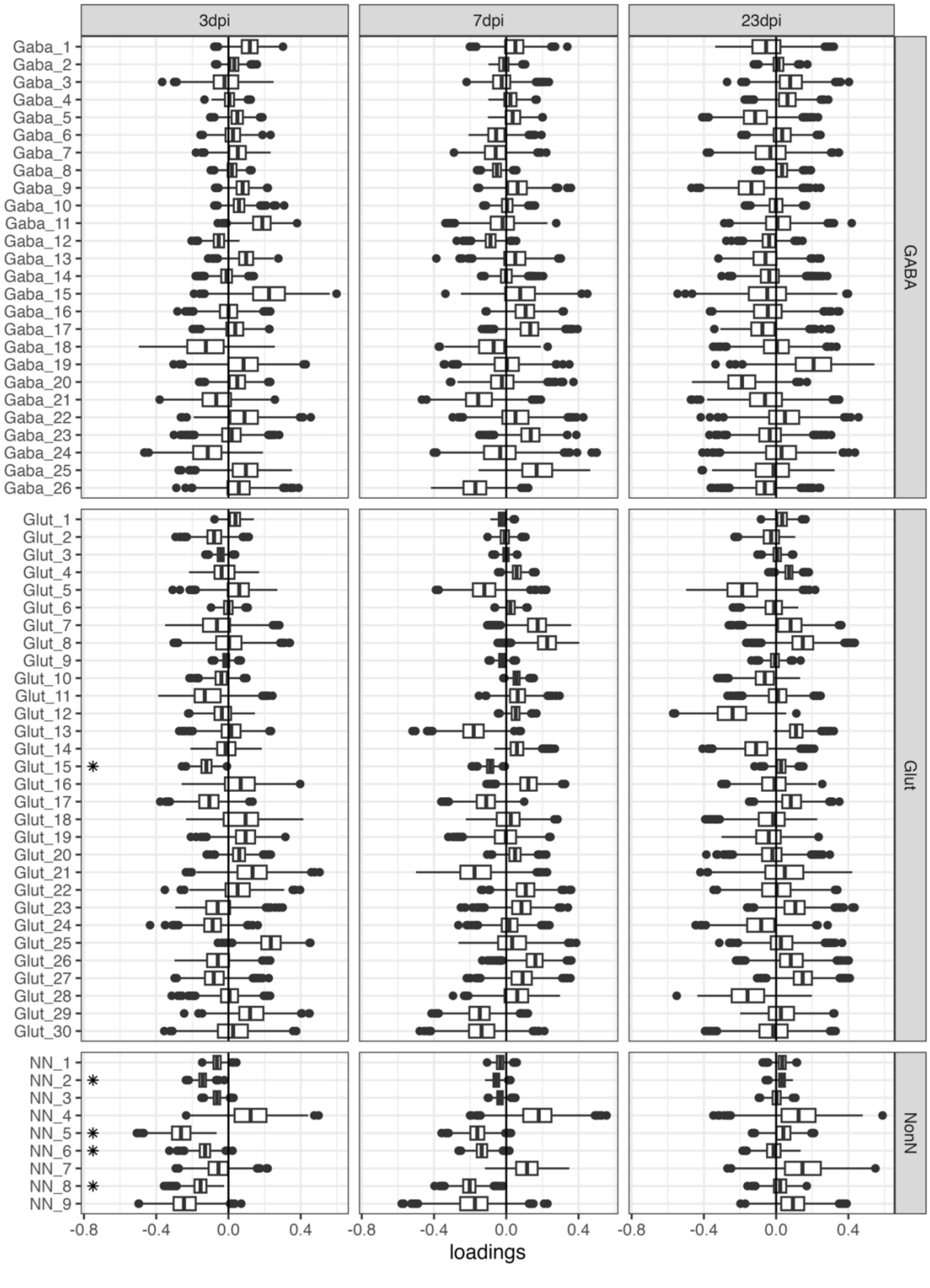


Supplementary Figure 15 **Cluster-based composition shifts calculated by Cacoa.** Box-plots shwoing cluster-based expression shifts of the different infected groups in comparison to the control calculated with the Cacoa package (Petukhov et al., 2022). Shifts on the x-axis in the negative space show a increase of cell proportions in the control samples, whereas a shift towards the positive space indicate an increase in the different infected groups (3, 7 and 23 dpi). Stars depict significance change in cell densities (P <= 0.05 after BH correction).

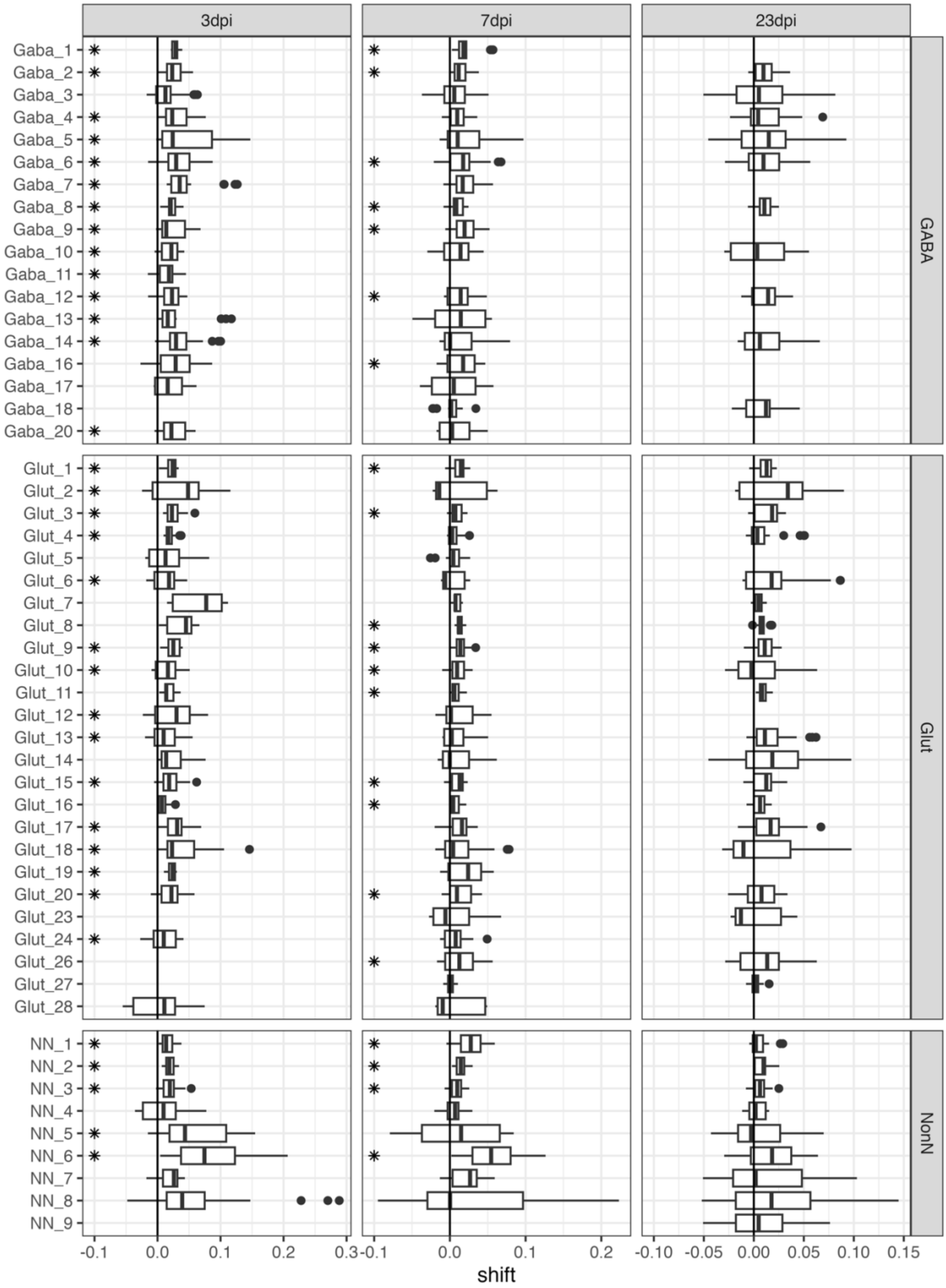


Supplementary Figure 16 **Cluster-based analysis of changes in expression magnitudes.** Boxplots show the shifts in expression magnitudes based on Cacoa package (Petukhov et al., 2022) in the different infected samples compared to the control group. Stars depict a significant change in expression in a cluster (P <= 0.05, after BH correction).

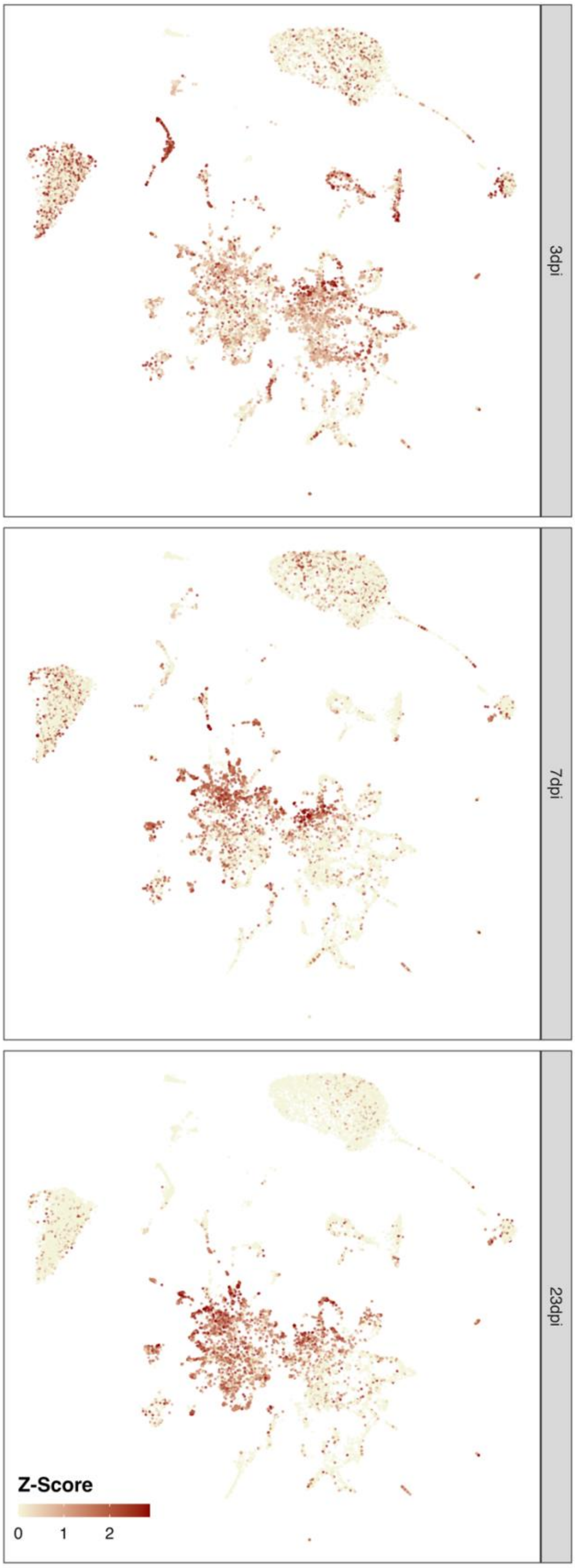


Supplementary Figure 17 **Cluster-free expression shifts.** UMAP embedding showing of the whole dataset showing adjusted statistical significance levels (color) of expression shift magnitudes. Analysis was performed based on Cacoa package (Petukhov et al., 2022).

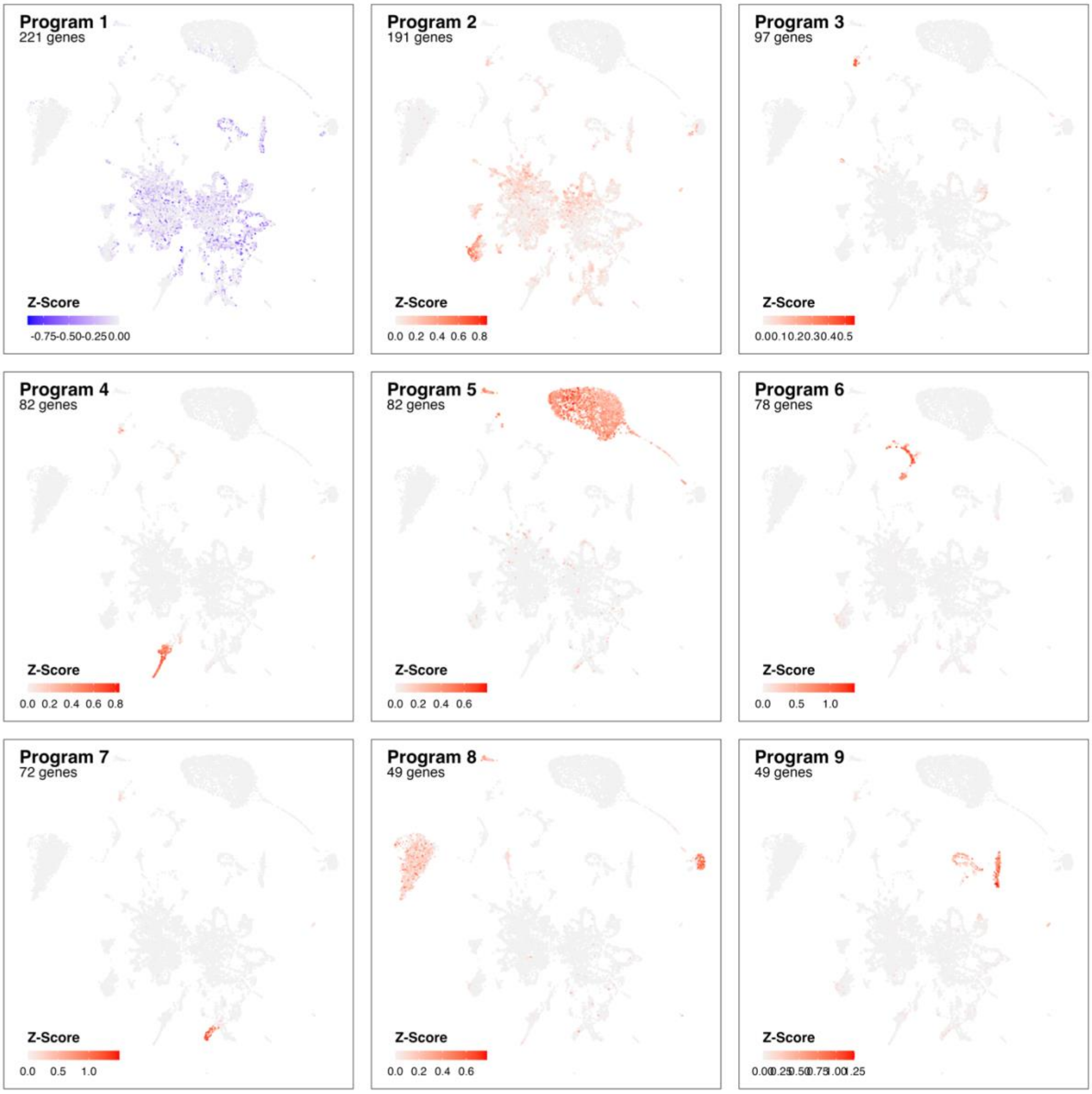


Supplementary Figure 18 **Identified gene programs at 3 dpi based on cluster-free genes expression analysis.** Adjusted z-scores are shown (color) for most pronounce genes programs identified comparing Control samples with infection at 3 dpi.

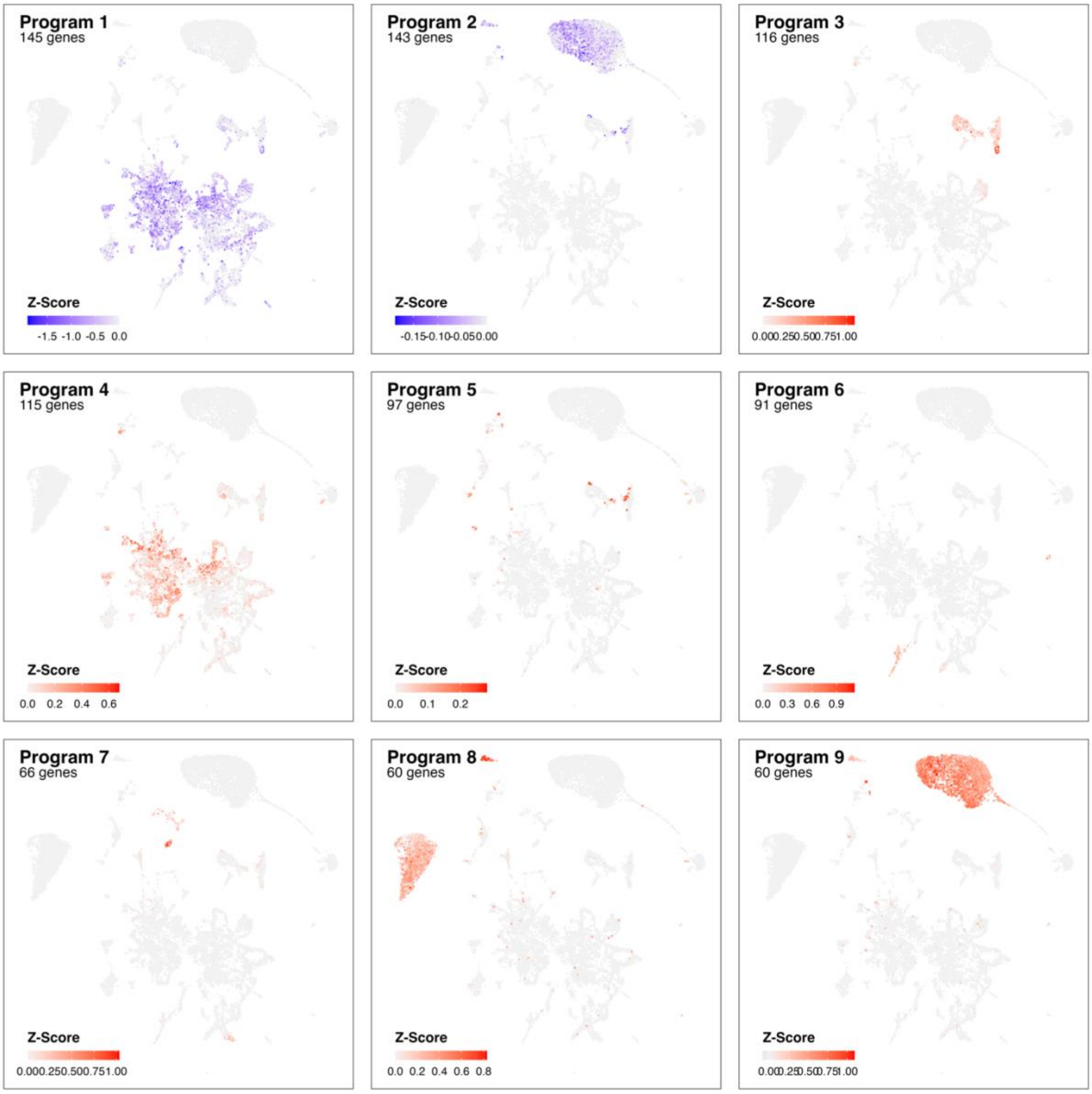


Supplementary Figure 19 **Identified gene programs at 3 dpi based on cluster-free genes expression analysis.** Adjusted z-scores are shown (color) for most pronounce genes programs identified comparing Control samples with infection at 7 dpi.

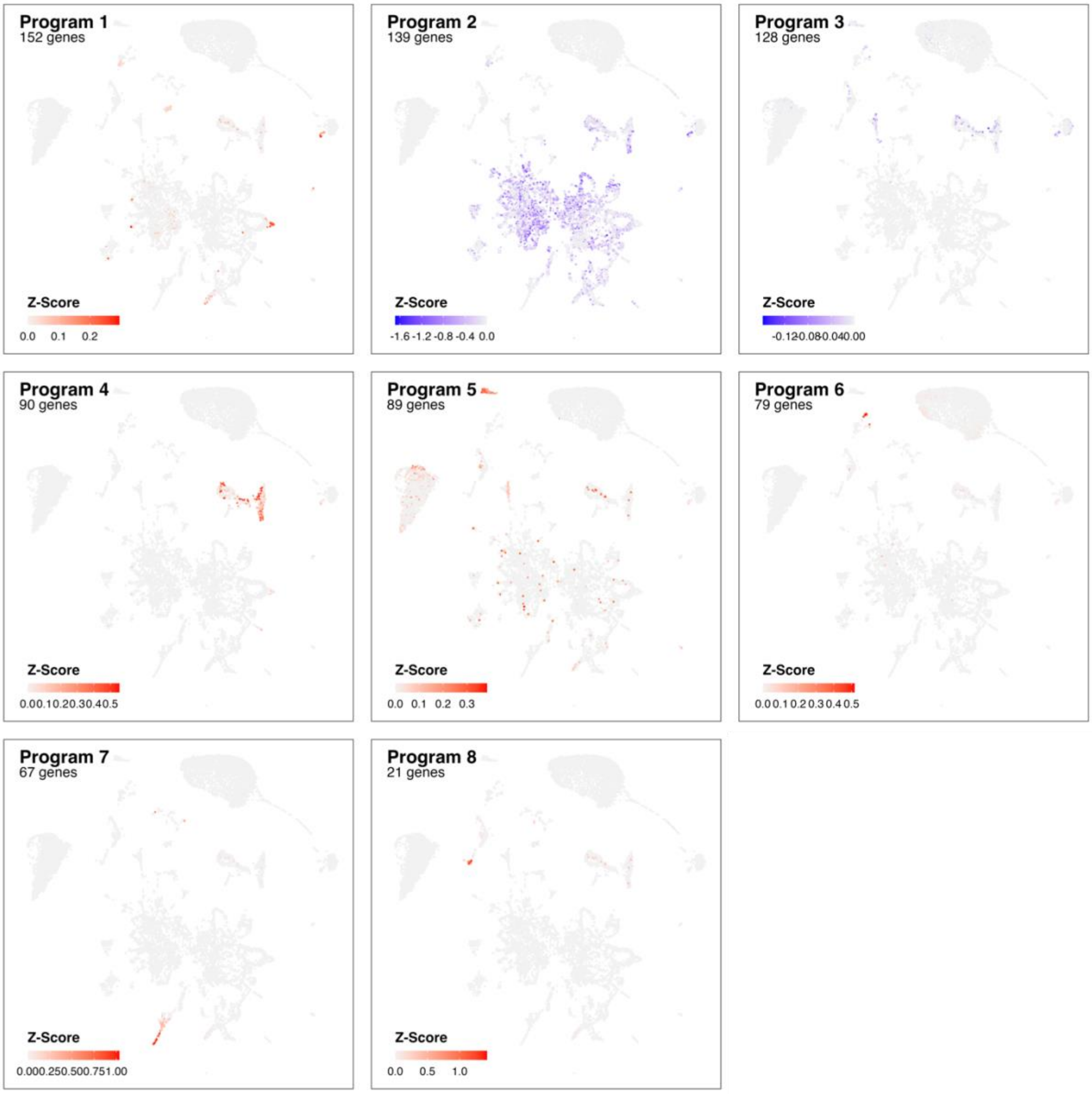


Supplementary Figure 20 **Identified gene programs at 3 dpi based on cluster-free genes expression analysis.** Adjusted z-scores are shown (color) for most pronounce genes programs identified comparing Control samples with infection at 23 dpi.

### Referencens


Campbell, J. N., Macosko, E. Z., Fenselau, H., Pers, T. H., Lyubetskaya, A., Tenen, D., Goldman, M., Verstegen, A. M., Resch, J. M., McCarroll, S. A., Rosen, E. D., Lowell, B. B., & Tsai, L. T. (2017). A molecular census of arcuate hypothalamus and median eminence cell types. *Nat Neurosci*, *20*(3), 484-496. https://doi.org/10.1038/nn.4495

Chen, R., Wu, X., Jiang, L., & Zhang, Y. (2017). Single-Cell RNA-Seq Reveals Hypothalamic Cell Diversity. *Cell Rep*, *18*(13), 3227-3241. https://doi.org/10.1016/j.celrep.2017.03.004

Hochgerner, H., Zeisel, A., Lonnerberg, P., & Linnarsson, S. (2018). Conserved properties of dentate gyrus neurogenesis across postnatal development revealed by single-cell RNA sequencing. *Nat Neurosci*, *21*(2), 290-299. https://doi.org/10.1038/s41593-017-0056-2

Mickelsen, L. E., Bolisetty, M., Chimileski, B. R., Fujita, A., Beltrami, E. J., Costanzo, J. T., Naparstek, J. R., Robson, P., & Jackson, A. C. (2019). Single-cell transcriptomic analysis of the lateral hypothalamic area reveals molecularly distinct populations of inhibitory and excitatory neurons. *Nat Neurosci*, *22*(4), 642-656. https://doi.org/10.1038/s41593-019-0349-8

Mickelsen, L. E., Flynn, W. F., Springer, K., Wilson, L., Beltrami, E. J., Bolisetty, M., Robson, P., & Jackson, A. C. (2020). Cellular taxonomy and spatial organization of the murine ventral posterior hypothalamus. *Elife*, *9*. https://doi.org/10.7554/eLife.58901

Moffitt, J. R., Bambah-Mukku, D., Eichhorn, S. W., Vaughn, E., Shekhar, K., Perez, J. D., Rubinstein, N. D., Hao, J., Regev, A., Dulac, C., & Zhuang, X. (2018). Molecular, spatial, and functional single-cell profiling of the hypothalamic preoptic region. *Science*, *362*(6416). https://doi.org/10.1126/science.aau5324

Petukhov, V., Igolkina, A., Rydbirk, R., Mei, S., Christoffersen, L., Khodosevich, K., & Kharchenko, K. V. (2022). Case-control analysis of single-cell RNA-seq studies. *bioRxiv*, 2022.2003.2015.484475. https://doi.org/10.1101/2022.03.15.484475

Steuernagel, L., Lam, B. Y. H., Klemm, P., Dowsett, G. K. C., Bauder, C. A., Tadross, J. A., Hitschfeld, T. S., Del Rio Martin, A., Chen, W., de Solis, A. J., Fenselau, H., Davidsen, P., Cimino, I., Kohnke, S. N., Rimmington, D., Coll, A. P., Beyer, A., Yeo, G. S. H., & Bruning, J. C. (2022). HypoMap-a unified single-cell gene expression atlas of the murine hypothalamus. *Nat Metab*, *4*(10), 1402-1419. https://doi.org/10.1038/s42255-022-00657-y

Zeisel, A., Hochgerner, H., Lonnerberg, P., Johnsson, A., Memic, F., van der Zwan, J., Haring, M., Braun, E., Borm, L. E., La Manno, G., Codeluppi, S., Furlan, A., Lee, K., Skene, N., Harris, K. D., Hjerling-Leffler, J., Arenas, E., Ernfors, P., Marklund, U., & Linnarsson, S. (2018). Molecular Architecture of the Mouse Nervous System. *Cell*, *174*(4), 999-1014 e1022. https://doi.org/10.1016/j.cell.2018.06.021
